# A metagenomics approach to identification of eukaryotes in metazoan-associated microbiomes

**DOI:** 10.1101/2023.11.16.567323

**Authors:** Audra L. Crouch, Laine Monsey, Cameron Ramos, Matthew Z. Anderson

## Abstract

**Background:** Microbial eukaryotes are integral components of the microbiome where they shape community composition and ecological interactions. However, the abundance and diversity of eukaryotic species within the microbiome, the ‘eukaryome’, remains poorly defined. These deficiencies arise from unresolved technical limitations in recovering DNA from microbial eukaryotes due to their relatively low abundance in most samples and resilience to extraction. To overcome these limitations, we developed an extraction protocol that specifically targets recovery of eukaryotic microbes from microbiome samples and allows for metagenomics sequencing of eukaryotic species.

**Methods:** Microbes were seeded in synthetic stool prior to DNA extraction to mimic microbiome samples from the gastrointestinal tract. Assessment of cell disruption was performed using intracellular staining with the azo dye trypan blue or quantification of DNA recovery. A mock microbial community of five bacteria and five eukaryotes was built to test the effectiveness of the full protocol by seeding stool with defined numbers of cells from each species.

**Results:** Mechanical disruption efficiently released DNA from bacterial, fungal, and protozoan species where standard microbiome DNA extraction kits did not. Optimization of the bead beating parameters lysed >95% of fungal cells within synthetic stool samples. In a mixed bacteria-eukaryote sample, eukaryotic DNA could be further enriched by targeting methylated DNA for destruction with methyl-specific restriction endonucleases. Application of this approach to a defined community of 10 different microbes, five eukaryotes and five bacteria, seeded in synthetic stool demonstrated the success of this strategy by enriching for eukaryotes approximately 72-fold and producing a eukaryote-dominated DNA pool.

**Conclusions:** Overall, development of a microbiome sample protocol that includes DNA extraction and enrichment from eukaryotic species will facilitate exploration of the eukaryome and its impact on human health.

## BACKGROUND

Microbial communities are globally ubiquitous entities that shape the biotic and abiotic makeup of their local environment. Constituents of these microbial communities, which include viruses, bacteria, archaea, and eukaryotes, mold their niche by forging complex ecological interactions among species within and across kingdoms (Robinson et al. 2010; Faust and Raes 2012; Medlock et al. 2018). In association with multicellular organisms from plants to animals, they modify the host environment including contributions to host development and physiology (Mazmanian et al. 2005; Bouskra et al. 2008; Ortiz-Castro et al. 2009; Marchesi et al. 2016). One of the most well-characterized microbial niches is the human gut, where the bacterial repertoire and their dynamic responses across a range of physiological states have been thoroughly examined (Turnbaugh et al. 2007; Lozupone et al. 2012; Yatsunenko et al. 2012).

These studies have revealed important causal links between individual microbes or community shifts, and clinical disease within the host including inflammatory bowel disease (IBD) (Miyoshi et al. 2018; Limon et al. 2019; Rangan et al. 2019), colorectal cancer (Coker et al. 2019; Koliarakis et al. 2019), and ankylosing spondylitis (Li et al. 2019). Furthermore, use of some bacteria as probiotics to alleviate human disease demonstrates the ability to reconfigure community composition to alter host health outcomes (Suez et al. 2018).

Despite these advances, the eukaryotic components of the microbiome, the ‘eukaryome’, have been largely unexplored and its consequences on human health and disease therefore remain mostly undefined (Gill et al. 2006; Parfrey et al. 2011; Huseyin et al. 2017).

Fungi, protozoans, and helminths have all been found within the human gastrointestinal tract (GIT) (Ghannoum et al. 2010; Hamad et al. 2014; Hugon et al. 2017), but typically comprise ≤1% of the microbial DNA recovered from fecal samples (Nam et al. 2008; Nash et al. 2017). This low abundance is further complicated by a lack of species and protocol consistency across studies that could conflate identification of residents of the human gut with environmental contaminants (Hamad et al. 2016), raising concerns that current definitions of the human gut eukaryome are particularly limited (Auchtung et al. 2018). Furthermore, descriptions of the eukaryome have commonly relied on industrialized populations that are associated with reduced taxonomic diversity and eukaryote abundance, thereby reducing the effectiveness of these efforts (Smits et al. 2017; Sonnenberg et al. 2019). Yet, fungi such as *Candida*, *Malassezia*, and *Saccharomyces* and the protist *Blastocystis* are commonly found across studies and stand as signature eukaryotes of the industrialized gut microbiome (Nash et al. 2017; Kodio et al. 2019; Limon et al. 2019)

Microbial eukaryotes play important and often unique ecological roles within microbial communities. These organisms are frequently an order of magnitude larger than their bacterial counterparts and therefore contribute considerably more to the microbial biomass and interactions that shape the gut niche than accounted for through sequencing (Baron 1996; Yang et al. 2011; Levin and Angert 2015). In fact, previous studies demonstrated that the abundance of specific microbial eukaryotes are associated with changes in bacterial composition (Audebert et al. 2016; Ramanan et al. 2016; Sam et al. 2017) and contribute directly to the spatial organization of bacterial species within various niches (Morton et al. 2015; Graham et al. 2017; Jenkins et al. 2017; Zhang et al. 2018). Some interactions are direct such as the secretion of surface-linked proteins by *Candida* species to modulate formation of mixed-kingdom biofilms (Klotz et al. 2007; Shirtliff et al. 2009). Other interactions occur by interfacing with host immunity (Parfrey et al. 2011; Limon et al. 2019). Overgrowth of *Candida albicans* in the gut leads to immune cell activation, production of inflammatory cytokines, and, ultimately, reduced bacterial alpha diversity (Iliev et al. 2012; Sokol et al. 2017). Similarly, secretion of the immunoregulatory glycoprotein ES-62 by the nematode *Acanthocheilonema vitae* suppresses

IL-17-mediated inflammation and overgrowth of commensal bacteria including Firmicutes and Proteobacteria (Smallwood et al. 2017; Doonan et al. 2019). Thus, in the absence of knowing the repertoire of gut-associated eukaryotes, our understanding of the full ecological context of the microbiome and the roles of community members within that niche is lacking.

A major hurdle to defining and studying the eukaryome has been the lack of consistent methodologies employed in species identification and quantification. Early analysis relied on successful culturing or microscopic identification of eukaryotes from fecal samples (Hsieh et al. 1965; Randremanana et al. 2012; Gouba et al. 2013), but has more recently been replaced with amplicon-based sequencing approaches. This technique utilizes nucleotide variants within the 18S ribosomal RNA gene and internal transcribed spacer region of the rDNA locus (e.g., *ITS1* and *ITS2*) to differentiate taxa and quantify their relative abundance at the genus- or species-level (Hoffmann et al. 2013; Nash et al. 2017). However, amplicon-based profiling introduces bias when assessing species composition in complex communities (Kim et al. 2017; Nash et al. 2017). The commonly used “pan-fungal” primers were designed based on alignment to a handful of pilot species and have an estimated taxonomic coverage of only ∼44% of fungal species known to be associated with mammalian niches based on microscopic and culture- based approaches (Kumar et al. 2015; Usyk et al. 2017). Furthermore, use of two different loci in taxonomic determination, *ITS1* and 18S rRNA gene, from the same human stool samples resulted in substantially different estimates of associated fungal species, highlighting the inadequacies of this approach (Nash et al. 2017). Additional limitations exist to commonly used methods to lyse cells in microbiome studies, which focus on bacterial and archaeal species and therefore are unable to break open the rigid cell walls and/or thick polysaccharide capsules surrounding many fungi (Doering 2009). As a result, metagenomics approaches from the Human Microbiome Project using whole genome sequencing (WGS) found only ∼0.0004% of all reads mapped to eukaryotic microbes (Nash et al. 2017). This lack of eukaryotic representation is acutely felt for gut-associated protozoans and helminths that are often entirely unreported (Hamad et al. 2016). Thus, a viable unbiased approach to identify eukaryotes within host- associated microbiomes is needed.

Here, we developed and tested a metagenomics approach for recovering eukaryotic species from representative fungi and protozoa seeded in synthetic stool. The developed method vastly outperformed all tested commercial microbiome kits for DNA extraction and recovery from microbial eukaryotes. Additional steps to enrich for eukaryotic material through methyl-specific DNA digestion further increased the relative recovery of eukaryotic DNA when mixed with bacterial cells. Application of the full protocol to a mock community of ten microbes heavily biased toward bacteria, led to a 70-fold enrichment of the eukaryotes and recovery of a eukaryote-dominant DNA pool. Thus, this approach to microbiome profiling now allows unbiased identification of microbial eukaryotes within stool samples and will greatly expand our understanding of the ecology of host niches.

## RESULTS

### DNA extraction of eukaryotes from fecal matter

A major hurdle to defining the human gut eukaryome has been the poor recovery of microbial eukaryote DNA from microbiome samples. To determine the efficacy of recovering eukaryotic DNA from stool samples, microbe-free synthetic stool was seeded with 1.0x10^8^ *Cryptococcus neoformans* cells per milliliter (mL), which is a particularly recalcitrant fungal organism to DNA extractions due to the presence of a tightly cross-linked cell wall and thick capsule of exoglycoprotein (O’Meara and Alspaugh 2012). Equal aliquots of synthetic stool seeded with *C. neoformans* were prepared for DNA extraction using three common microbiome preparation kits, a fungal-specific DNA isolation kit, and mechanical lysis via bead beating.

Either bead beating or the fungal-specific kit efficiently recovered DNA from *C. neoformans*; whereas, all three microbiome kits resulted in significantly reduced DNA yields (Figure 1, one- way ANOVA (F(4,15) = 36.0, p < 1.0E-4). Thus, we pursued a eukaryome extraction approach based on mechanical disruption via bead beating as it significantly outperformed standard microbiome extraction protocols and could be easily modified for optimization.

**Figure 1.**
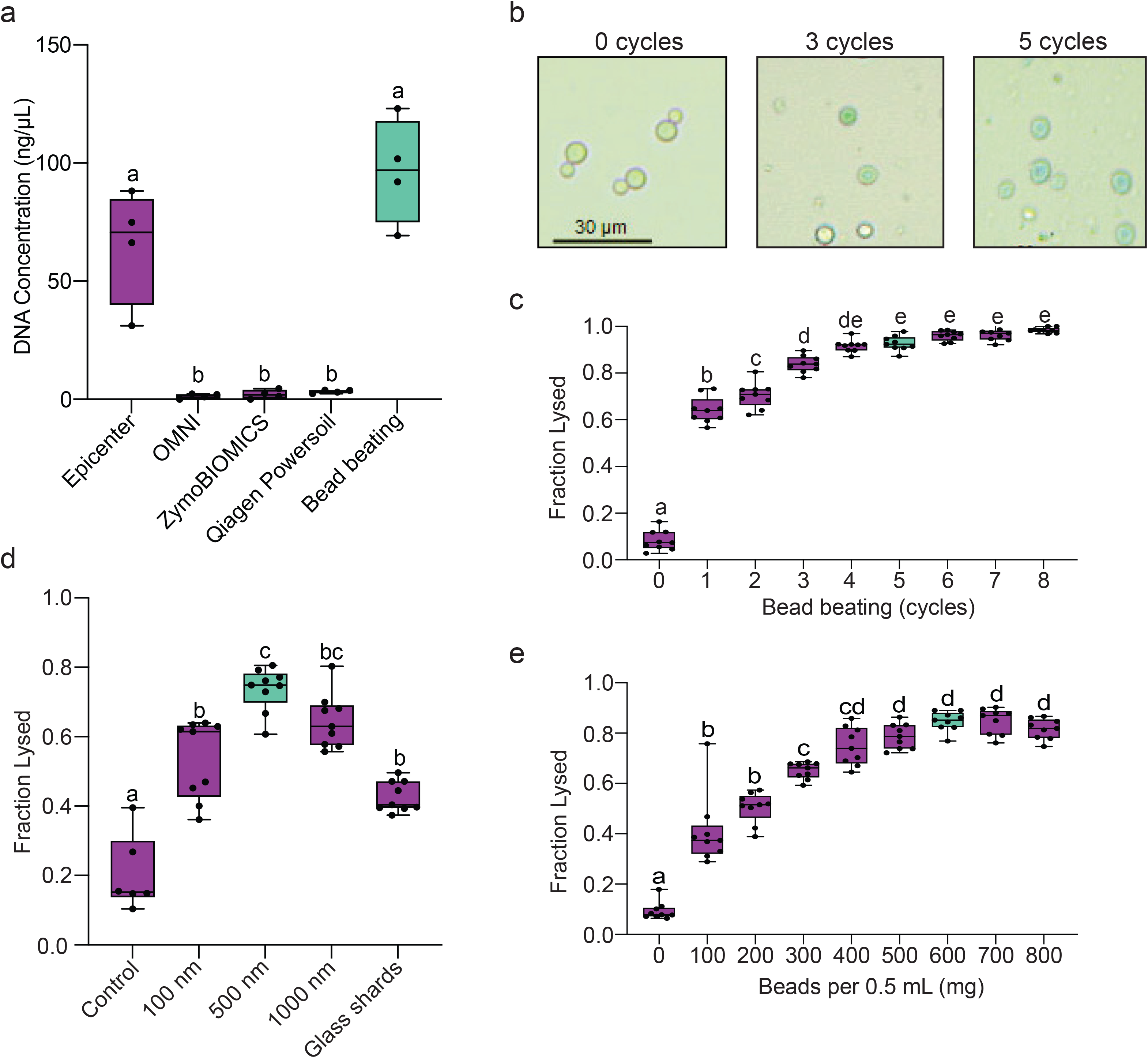
Bead beating effectively recovers DNA from fungal cells. Synthetic stool (500 mg) was seeded with 1x10^8^ *C. neoformans* cells and the efficiency of five different DNA recovery approaches for microbiome analysis was tested. The amount of DNA recovered using each approach was plotted where boxplots mark the interquartile ranges with whiskers extending to the outermost points. Letters indicate statistically different approaches based on a one-way ANOVA and Tukey’s posthoc test at p < 0.05. N = 5.

To determine the optimal conditions that efficiently release DNA during bead beating, we incrementally adjusted individual parameters and assessed their effect on maximizing cell lysis. Visual identification of lysed versus intact *C. neoformans* cells can be challenging so the azo dye trypan blue was added to distinguish between lysed cells, which appear blue, and intact cells that lack staining (Figure 2A). First, the effect of increasing the number of bead beating cycles (one-minute on, two-minutes off) on cell lysis was assessed. Cell lysis increased with added cycles, reaching a peak effectiveness beginning at 5 cycles (Figure 2A, B). Additional cycles did not increase cell lysis and led to further cellular fragmentation. Next, beads of various sizes ranging from 100 to 1,000 nanometers (nm) were tested for cell disruption alongside glass shards and no beads, which served as a negative control. Beads averaging 500 nm in diameter proved most effective at lysing *C. neoformans* cells compared to other bead sizes and glass shards (Figure 2C). Finally, the effect of the amount of beads per 0.5 mL of *C. neoformans* sample was assayed. Cell lysis increased with added beads up to 600 milligrams (mg) and decreased slightly with greater amounts (Figure 2D). Thus, lysis of *C. neoformans* was most effective using 600 mg of 500 nm glass beads for a duration of five cycles of bead beating.

**Figure 2.**
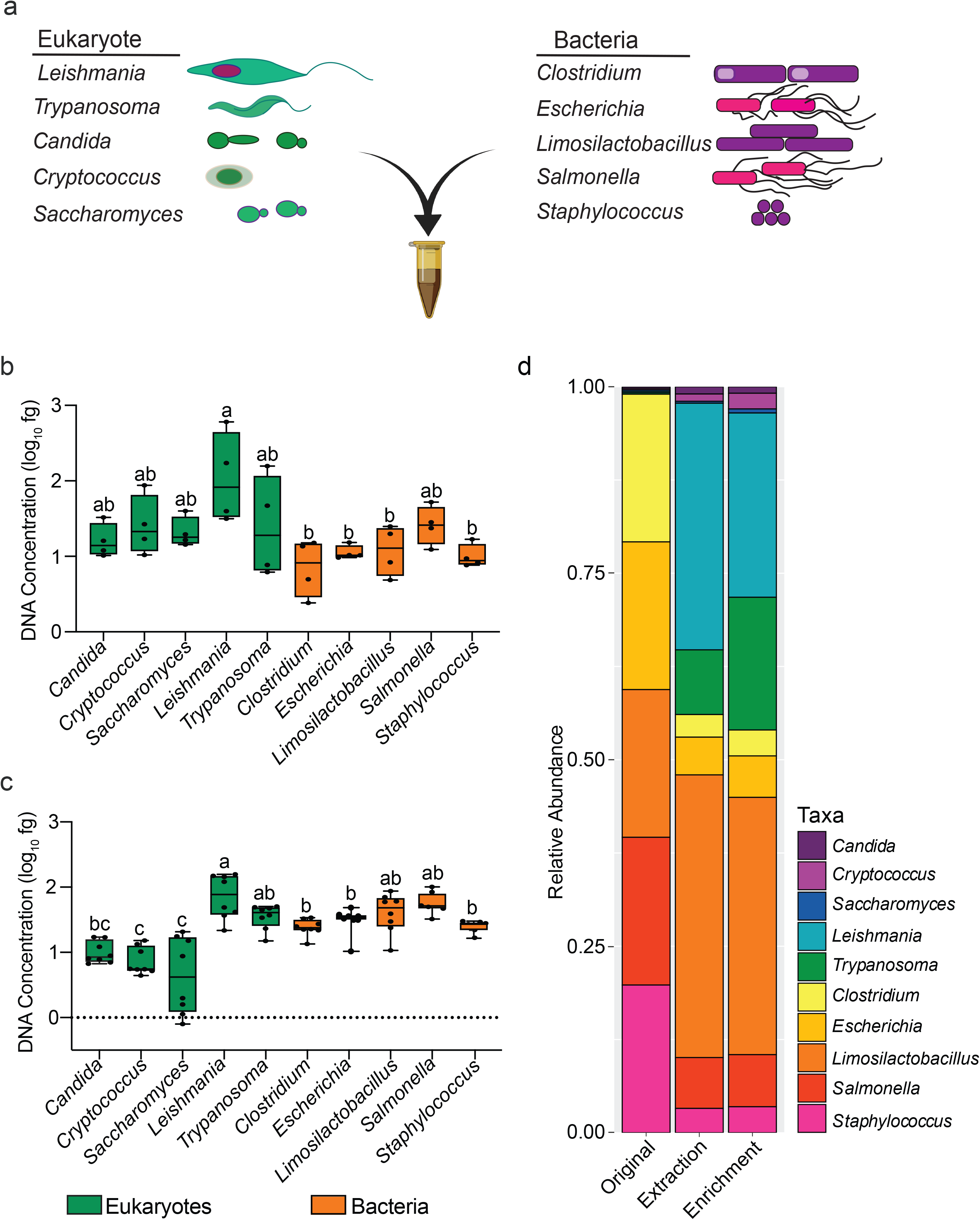
Bead beating optimization for maximal fungal cell lysis. A. *C. neoformans* cells (1x10^8^) were bead beat with 100 μm beads for different cycles (one minute on, 2 minutes off) and incubated with 0.4% trypan blue solution. Intact cells appear clear but lysed cells stain blue. Scale bar = 30 μm. The fraction of *C. neoformans* cells lysed as indicated by trypan blue staining was measured across a range of bead beating cycles with 100 μm glass beads (**B**), glass bead sizes and glass shards for 5 cycles (**C**), and mass of 100 μm glass beads per 0.5 mL of sample for 5 cycles (**D**). Six fields of view were analyzed per replicate. Boxplots mark the interquartile ranges with whiskers extending to the outermost points. Letters indicate statistically different fractions of cell lysis based on a one-way ANOVA and Tukey’s posthoc test at p < 0.05. N = 9.

The buffer used to homogenize stool samples is also critical to cell lysis and DNA recovery (Huseyin et al. 2017; Frau et al. 2019). To test the effect of commonly used lysis buffers on DNA recovery, we included buffers from microbiome extraction kits, the commonly used extraction buffer cetyltrimethylammonium bromide (CTAB), and a buffered detergent solution (TENT) with or without 3% sodium dodecyl sulfate (SDS; TENTS). Surprisingly, CTAB and kit-associated buffers performed poorly compared to TENT and TENTS buffer during bead beating, although the SDS in TENTS buffer also reduced cell lysis and DNA recovery (Figure 3A).

**Figure 3.**
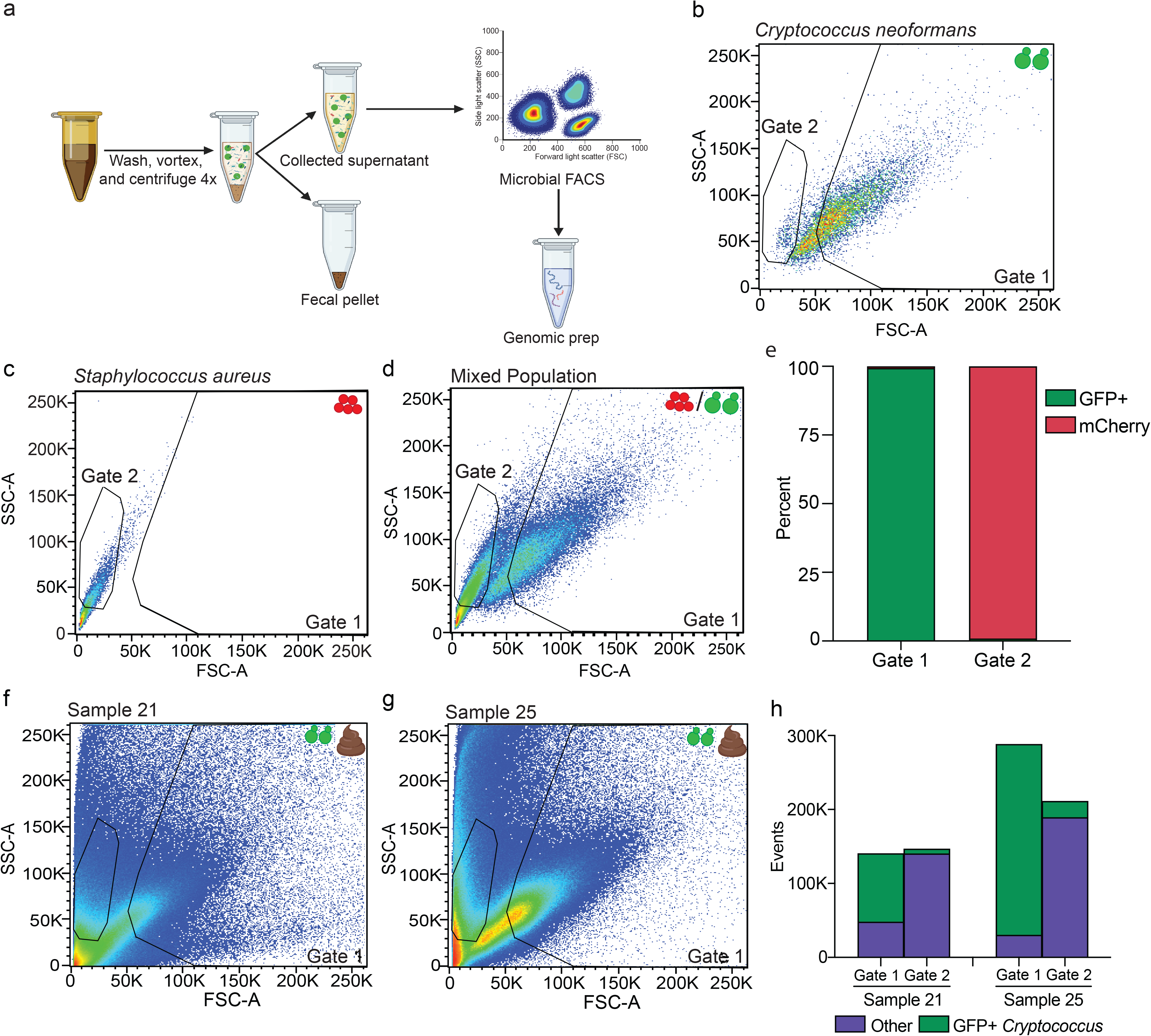
Optimization of DNA recovery from fungal cells. A. *C. neoformans* cells seeded in synthetic stool (1x10^8^ cells in 500 mg) were suspended in 500 uL of various buffers commonly used in DNA extraction from microbiome samples. DNA recovery was plotted for each buffer. Boxplots mark the interquartile ranges with whiskers extending to the outermost points. Letters indicate statistically statistically different DNA yields based on a one-way ANOVA and Tukey’s posthoc test at p < 0.05. N = 9. **B.** DNA was extracted from synthetic stool as in **A** from different volumes of synthetic stool in TENT buffer seeded with the same density of *C. neoformans* cells. The percentage of maximal DNA that could be recovered from 1x10^8^ cells in 500 mg of stool is plotted when DNA was extracted and purified with either phenol:chloroform (red), phenol:chloroform and an additional chloroform purification (blue), or phenol:chloroform, an additional chloroform purification, and magnetic bead purification (green). The black line indicates the maximal theoretical recovery across different initial masses of stool. Data is plotted as averages with standard deviation. N = 3

Unexpectedly, quantification of DNA yields from *C. neoformans* in synthetic stool did not initially correlate with increasing numbers of seeded cells, suggesting carryover of contaminating materials from the synthetic stool inhibited accurate quantification of recovered DNA. However, addition of magnetic bead purification to purify DNA following phenol-chloroform and a second chloroform extraction removed the contaminating substance(s) and led to a strong association between the seeded number of cells and DNA recovery (cor = 0.984, Pearson’s test = 10.99, df = 4, p-value = 1.00E-3, Figure 3B).

### Enrichment of eukaryotic DNA from stool

Another major hurdle to studying eukaryotes within metazoan-associated gut microbiomes is the overwhelming presence of bacteria and archaea within these samples (Frau et al. 2019). To improve identification of eukaryotes within metagenomics samples, we exploited fundamental differences in methylation states of bacterial and eukaryote genomes. Bacterial DNA is commonly methylated at the N-6 position of adenosine (^m6^A) and C-5 or N-4 position of cytosine residues; whereas eukaryotes typically lack these modifications, allowing their genomes to be distinguished through the use of methyl-specific restriction endonucleases. For example, the DpnI restriction enzyme recognizes ^m6^A and cleaves the sequence 5’-G^m6^ATC- 3’ that is present in bacterial but rarely eukaryotic genomes. To assess the specificity of methyl-specific restriction enzymes towards digesting bacterial genomes, we incubated DNA from the Gram-negative bacterium *Escherichia coli* and *C. neoformans* with DpnI either alone or mixed together in equivalent ratios. DpnI rapidly degraded *E. coli* DNA, reducing large DNA molecules to small fragments, but left *C. neoformans* DNA largely unaltered (Figure 4a). Quantitative PCR (qPCR) of individual loci encoding a DpnI recognition sequence from either species in a 50:50 mix produced a five-fold reduction in intact *E. coli* but not *C. neoformans* DNA (Figure 4B, S1). Although DpnI fragments the genome, it does not reduce DNA to free nucleotides.

**Figure 4.**
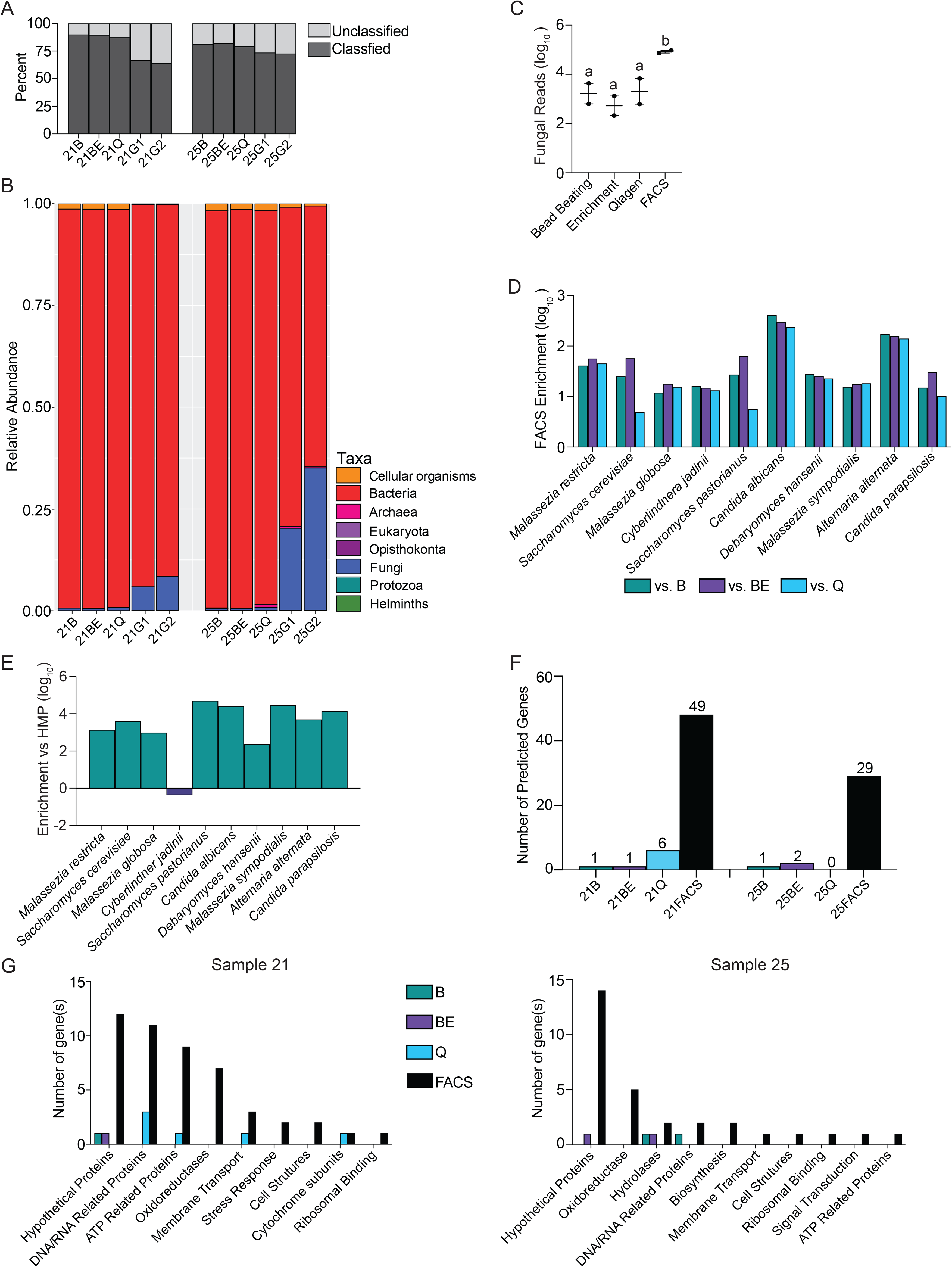
Methyl-sensitive restriction enzymes digest bacterial but not eukaryotic DNA. A. Two 1 μg aliquots of *E. coli* (left), *C. neoformans* (right), or a 50:50 mix of *E. coli*:*C. neoformans* (center) DNA was incubated with the DpnI restriction enzyme or mock treated for 1 hour and run by gel electrophoresis. DNA sizes are marked by a 10 kb ladder. **B.** 50:50 mixes of *E. coli* and *C. neoformans* DNA was digested with DpnI or mock treated for 3 hours and the fraction of input DNA recovered after purifying each reaction with 0.7x magnetic beads. **C.** DNA was digested as in **B** and purified with varying ratios of magnetic beads compared to the reaction volume. The fraction of recovered DNA compared to the input amount is plotted as the average and standard deviation. N = 3.

Consequently, the method of DNA purification following digestion can modulate the recovery of eukaryotic and bacterial DNA. Use of bead-to-sample volume ratios below 0.7x failed to recover all *C. neoformans* DNA and did not significantly decrease the amount of recovered *E. coli* DNA (Figure 4C). Thus, DpnI digestion can selectively enrich for eukaryotic DNA within a mixed pool by degrading bacterial genomes that are then selectively lost during DNA recovery.

### Eukaryotic DNA enrichment from a defined microbial community

To test the efficacy of the developed protocol with more representative microbial communities, we built a ten-member mock community consisting of five eukaryotes (*Candida albicans*, *C. neoformans*, *Saccharomyces cerevisiae*, *Leishmania major*, and *Trypanosoma brucei*) and 5 bacterial species (*Clostridioides difficile*, *E. coli*, *Lactobacillus reuteri*, *Salmonella typhimurium*, and *Staphylococcus aureus*) (Figure 5A). The mock communities were seeded in synthetic stool at either equivalent ratios (1:1) or a 99:1 (bacteria:eukaryote) ratio of cells. We extracted DNA from the mock community using the described bead beating protocol and quantified abundance of each species by qPCR for a set of species-specific, single-copy loci for each microbe (Table 1). This approach recovered similar amounts of DNA from each species when mock communities were seeded with equivalent numbers of cells from each organism, although a bias for recovering *L. major* DNA was found (one way ANOVA, F(9,30) = 3.078, p=0.0098, Figure 5B). If DNA extraction from bacteria and eukaryotes is also similar in the 99:1 bacterial-dominated mock community, bacterial DNA should be present in near 100-fold excess of eukaryotic DNA. In contrast, DNA from some eukaryotes was only ten-fold less than DNA from bacterial species (*C. albicans, C. neoformans*, and *S. cerevisiae*) and, surprisingly, *L. major* and *T. brucei* were present in equal concentration to seeded bacteria, indicating an enrichment for eukaryotes from bead beating alone (one-way ANOVA, F(9, 70) = 17.47, p<0.0001, Figure 5C). This bias resulted in a 52x enrichment of eukaryotic DNA compared to their initial cellular representation within the defined community (Figure 5D).

**Figure 5.**
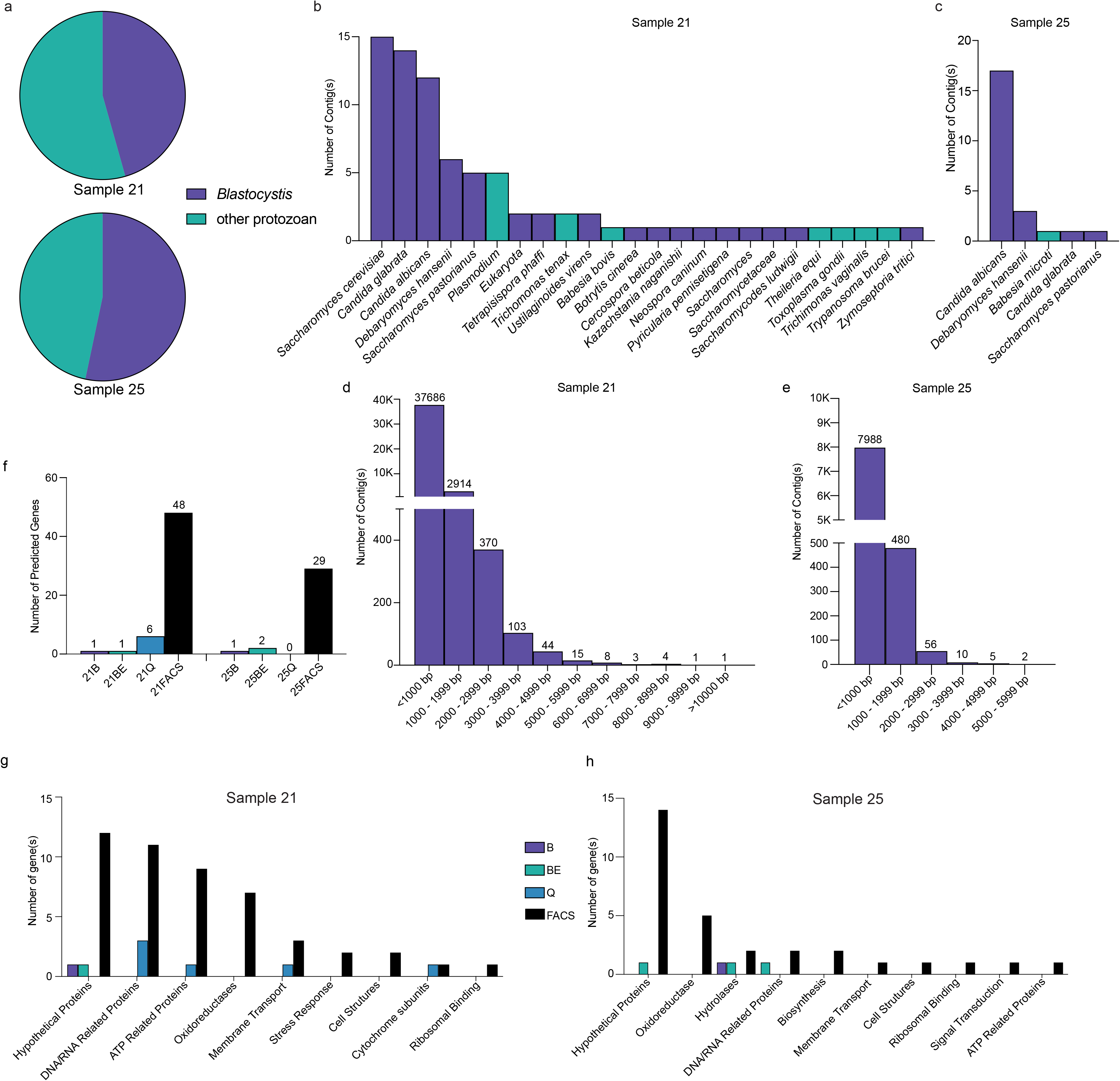
A eukaryome enrichment protocol preferentially recovers eukaryotic DNA. A. Five bacterial and five eukaryotic species were seeded in 500 mg of synthetic stool as either equivalent (1:1) or bacterial-biased (99:1) ratios with equal representation of species within the bacterial or eukaryotic groups. The developed eukaryome protocol was performed on either 1:1 (B) or 99:1 (C) microbial communities and the fraction of DNA recovered from each species plotted. Recovered DNA was determined using quantitative PCR with species-specific primers against a single genomic locus in each organism and quantified using a standard curve of known DNA concentrations. Boxplots mark the interquartile ranges with whiskers extending to the outermost points. Letters indicate statistically different DNA recovery based on a one-way ANOVA and Tukey’s posthoc test at p < 0.05. N = 4. (D). DNA recovered from 99:1 mixed communities was treated with the DpnI restriction enzyme for 3 hours and the fraction of DNA recovered following purification with 0.7x magnetic beads determined by qPCR and plotted for nine community members A solid line indicates the mean and whiskers indicate the standard deviation. E. The amount of DNA recovered from preparation of the 99:1 community is shown for L. reuteri either before (extracted) or after (enriched) treatment with the DpnI restriction enzyme for 3 h. F. The relative abundance of DNA quantified by qPCR of single locus genes using species-specific primers is given for the initial community (left), following DNA extraction (middle), or after enrichment by DpnI digestion (right). Each species is represented as the average of four biological replicates and color coded. L. reuteri was removed from the enrichment column due to a lack of DpnI recognition site within the diagnostic amplicon.

DNA harvested from the 99:1 bacteria-dominated samples was subjected to eukaryote enrichment using the methyl-specific DpnI restriction enzyme. Following digestion for three hours, DNA was recovered, and species abundance re-assessed via species-specific primers.

Enzymatic digestion reduced the recovered DNA from all bacterial species encoding a DpnI recognition site within the diagnostic locus by approximately 4-fold (Figure 5E). *L. reuteri* DNA was not depleted as it lacks a DpnI site within the gene used to measure abundance (Figure 5F, Figure S1). DNA from eukaryotes was generally unaffected by DpnI digestion with the exception of *C. albicans* and *L. major* that underwent a four-fold reduction in DNA recovery (Figure 5E).

This resulted in a roughly 2-fold enrichment for eukaryote DNA compared to the input. Taken together, the eukaryome protocol developed here enriched for eukaryotes by 73-fold (Figure 5D), shifting eukaryotic representation from a scarce number of representative cells to the dominant product in final DNA pools.

## CONCLUSIONS

Technical limitations and low representation in microbiome samples have impeded investigation of the microbial eukaryotes residing within the human gastrointestinal tract. Our approach used mechanical disruption by bead beating to lyse resilient eukaryotic cells followed by digestion of methylated DNA to selectively enrich for eukaryotic DNA from mixed microbial communities. Testing the technique with a synthetic microbial community produced a 70-fold enrichment of eukaryotic DNA that would skew metagenomic sequencing of microbes in canonical stool samples towards eukaryotes to facilitate unbiased identification of eukaryome composition across treatments and populations. Thus, this approach overcomes constraints in releasing eukaryotic DNA and recovering it from microbiome samples dominated by bacteria and archaea.

This protocol addresses systematic bias found in amplicon-based community profiling of microbial eukaryotes in metazoan-associated microbiomes. Thorough comparative analysis between amplicon and metagenomics sequencing of bacterial and archaeal communities revealed limitations based on sequencing errors, amplification biases, and the inability to access gene content information (Shakya et al. 2013; Poretsky et al. 2014). As a result, most microbiome studies have shifted towards metagenomic approaches. Yet, amplicon-based sequencing remains the primary method to determine the taxonomic distribution of resident eukaryotes. Our approach now allows for the application of metagenomic sequencing to identify eukaryotes within microbiome samples, which will remove similar biases in taxonomic identification and allow for gene-level analysis of recovered eukaryotes. Slight variation in recovery of different microbial eukaryotes occurred in both the 1:1 and 99:1 communities, but DNA yields were within a much tighter range than what can be achieved with amplicon-based sequencing for many eukaryotic species (Nash et al. 2017). This vastly improved precision will facilitate comparisons of relative species abundance within samples and comparisons of the same species across populations or conditions to identify altered eukaryome states.

Furthermore, metagenomics approaches will yield the DNA sequences necessary to mine for altered gene functions that have proved particularly insightful in bacterial studies (Tasse et al. 2010; Almeida et al. 2019). Metagenome assembled genomes of eukaryote species remains a lofty ideal that has met with some success, but presently remains complicated by difficulties in correctly binning individual chromosomes into a single genome (West et al. 2018).

Mechanical lysis to extract DNA from microbial eukaryotes proved to be critical for eukaryotic enrichment. Bead beating has been a long-standing gold standard for cell lysis in fungi and other microbes (van Burik et al. 1998), and efficiently broke open highly resilient fungal cells and various protozoa. However, bead beating alone is not likely to have caused the drastic increase in recovered DNA as this step is present in some fecal sample kits. The two approaches with significantly improved DNA extraction did not include filtering steps that are common components of microbiome kits to separate DNA from cell debris. Thus, use of microbiome kits that include filtration may directly contribute to the unexpectedly low yields of eukaryotic DNA from metagenomics studies (Nash et al. 2017). Additionally, optimization of bead beating for eukaryotic recovery substantially increased DNA yields. We hypothesize that the larger cell size of eukaryotes made them more susceptible to being crushed by the larger glass beads whereas bacterial cells could easily slide between beads as they collided. This led to a 52x increase in eukaryotic DNA recovery compared to bacteria through bead beating alone (Figure 5). Decreased bead size, which performed less well for eukaryotes, may preferentially lyse bacterial cells and allow for profiling of bacteria and archaea within a sample that is split to allow simultaneous profiling of both populations. Collectively, these data suggest that kits should be generally avoided or substantially modified when planning to investigate microbial eukaryotes.

Differential digestion of methylated DNA only slightly favored eukaryotic enrichment. Targeting fundamental differences in methylation states between eukaryotes, bacteria, and archaea is a classic technique used for a variety of molecular biology techniques (Gruenbaum et al. 1981; Cross et al. 1994). Applied to the mixed microbial community here it had only a minor effect on further depleting bacterial DNA despite all species, except *L. reuteri*, encoding a DpnI cut site within the target gene used for quantification. Insufficient depletion could result from incomplete digestion although longer incubations and use of stationary phase cells, which should not contain large amounts of newly replicated and hemi-methylated DNA, did not increase the efficiency of removing bacterial DNA. This suggests that residual bacterial DNA remains despite digestion and selection of large DNA fragments via magnetic bead purification. This is unlikely to be a result of incomplete methylation of all ^m6^A sites as approximately 94% of all adenines available to methylation are modified in *E. coli* (Fang et al. 2012). Alternative approaches to extract DNA containing methylated nucleotides such as using a catalytically- inactive DpnI restriction enzyme (Barnes et al. 2014) could aid in further separating the eukaryotic DNA but become prohibitively expensive when working with large sample sets.

Importantly, ^m6^A is also present in eukaryotic genomes and could explain the reduction of *C. albicans* and *L. major* although neither species has specifically been shown to encode these modifications (Alderman and Xiao 2019). It is tempting to speculate that eukaryotic enrichment will be even more greatly enhanced in metagenomics sequencing by the large number of DpnI cut sites in the genomes of bacteria and archaea and larger genome sizes of microbial eukaryotes.

Ultimately, increasing access to study the eukaryome from microbiome samples facilitates pursuit of new research questions and the ability to revisit prior work. Questions remain as to the eukaryotic microbe repertoire of humans and other mammalian systems. Furthermore, their role in contributing to health and homeostasis remains poorly defined. This methodology allows not only the inclusion of the eukaryome in future studies but the ability to revisit this component of the microbiome in banked stool samples with the potential to produce major gains in our understanding of microbial ecology. Together, use of this approach and metagenomics sequencing can divulge the impact of the eukaryome in contributing to health and disease.

## METHODS

### Strains and growth conditions

Species used in this study are listed in Table 1. To produce cells for seeding synthetic stool, *C.albicans*, *C.neoformans*, and *S.cerevisiae* were grown in Yeast Peptone Dextrose (YPD; 20 g peptone, 10 g yeast extract, 50 mL of 40% glucose, and 2.5 mL of 10 mg/mL uridine in 1 L) medium overnight at 27°C. *L. major* was grown in 10 mL flasks in Modified M199 Medium (1- 560 mL consisting of 500 mL of M199, 50 mL of heat inactivated fetal bovine serum, 5 mL Penicillin /Streptomycin solution (10,000 U/10 mg per mL), and 5 mL of 1M HEPES solution) for 2-3 days at 27°C to reach full confluency. *T.brucei* was grown in 10 mL flasks with SDM-79 Medium (1-L medium consisting of 7 g MEM Powder, 2 g Grace’s Insect medium, 8 mL of 50x MEM essential amino acids, 6 mL of 100x MEM non-essential amino acids, 1 g glucose, 8 g HEPES, 5 g MOPS, 2 g sodium bicarbonate, 0.1 g sodium pyruvate, 200 mg DL-alanine, 100 mg L-arginine, 300 mg L-glutamine, 70 mg DL-methionine, 80 mg L-phenylalanine, 600 mg L-proline, 60 mg DL-serine, 160 mg taurine, 350 mg DL-threonine, 100 mg L-tyrosine, 10 mg guanosine, 4 mg folic acid, 50 mg D(+)glucosamine hydrochloride, 2 mg p-aminobenzoic acid, and 0.2 mg biotin) at 37°C for 2-3 days to reach full confluency.

For bacteria, *E.coli*, *S.aureus*, and *S.typhimurium* were grown in Luria-Bertani broth (LB) (1-L medium consisting of 10 g tryptone, 5 g yeast extract, and 10 g sodium chloride) overnight at 37°C. *L. reuteri* was grown in MRS broth (BD Difco) (1-L medium consisting 2 g dipotassium hydrogen phosphate, 20 g glucose, 0.2 g magnesium sulfate heptahydrate, 0.05 g manganous sulfate tetrahydrate, 8 gr meat extract, 10 g peptone, 5 g sodium acetate trihydrate, 2 g triammonium citrate, 4 yeast extract, pH 6.2) for 48 hours at 37°C. *C.difficile* was grown in Reinforced Clostridial Medium (RCM) (1-liter medium consisting of 3 g yeast extract, 10 g lab- lemco powder, 10 g peptone, 5 g glucose, 1 g soluble starch, 5 g sodium chloride, 3 g sodium acetate, 0.5 g cysteine hydrochloride) overnight at 37°C with an atmospheric gas mixture of 5% H_2_ - 15% CO_2_ - 85% N_2_.

### Assessment of fungal cell lysis

Overnight cultures of *C.neoformans* were counted via a hemocytometer and then diluted to 1.0 x 10^8^ cells/mL. Cells were centrifuged at 5000 rpm for 5 min, washed with 1 mL of 1x PBS, centrifuged at 5000 rpm for 5 min, and then resuspended in 1 mL of sterile synthetic stool. DNA extractions of *C.neoformans* inoculated synthetic stool samples were performed using the following kits: Epicenter Masterpure Yeast DNA Purification Kit, Omni Fecal DNA Purification Kit, ZymoBIOMICS DNA Extraction Kit, and Qiagin Powersoil Kit. All kits were used following guidelines and protocols specified by the manufacture. *C.neoformans* inundated synthetic stool samples were extracted using bead beating in TENT buffer (10 mM tris-HCl, 1 mM EDTA, 0.1 M sodium chloride, and 5% [v/v] triton X100, pH 8) for 5 minutes with 500 nm beads and purified as previously described (Suliman et al. 2007).

### DNA quantification

DNA quantification was determined fluorometrically using the Qubit 3.0 Fluorometer with the Qubit dsDNA High Sensitivity Assay Kit or the Qubit dsDNA Broad Range Assay kit (Thermo Fisher Scientific, Waltham, MA). DNA purity was determined via the 260/280 and 260/230 spectrometry ratios on a Nanodrop One (Thermo Fisher Scientific, Waltham, MA).

### Optimization of lysis buffer and contaminant removal

To determine optimal bead beating buffers, 10^8^ *C.neoformans* cells were resuspended in 500 uL of different lysis buffers in 2 mL threaded cryotubes: TENT (10 mM tris-HCl, 1 mM EDTA, 0.1 M sodium chloride, 5% [v/v] triton X100, pH 8), TENTS (TENT with 3% SDS), cetyl trimethyl ammonium bromide (CTAB), Qiagen Powersoil buffer (Qiagen, Hilden, Germany), and Epicenter Masterpure Yeast DNA Purification buffer (Epicentre, Madison, WI). Cells were bead beat for 5 cycles (1 minute on, 2 minutes off) using a BioSpec Mini-Beadbeater 8 (Biospec, Bartlesville, OK) with 0.6 g of 500 nm glass beads. The supernatant was removed follow mechanical lysis, added to two volumes of 95% ethanol and spun at 4°C for 15 minutes at 12000 rpm. The resulting pellet was washed with 70% ethanol and resuspended in 50 uL of water.

To determine the optimal sample mass for DNA extraction, 10^8^ *C.neoformans* cells were inoculated into 5 different masses (500, 250, 166, 125, and 100 mg) of sterile synthetic stool (Claremont BioSolutions, Upland, CA). Samples were bead beat in TENT buffer as described above and DNA recovered by phenol chloroform extraction of the supernatant. Briefly, 300 uL of Phenol:Chloroform:Isoamyl Alcohol (25:24:1) and 200 uL of 1x TE was added to the cryotube, and the tubes were rocked back and forth to fully homogenize the mixture. In a centrifuge cooled to 4°C, the cell-phenol mixture was centrifuged at 14,000 rpm or 10 min. DNA was precipitated and concentration and purity was assessed as described above. Following quantification, 400 uL of Chloroform:Isoamyl Alcohol (24:1) was added to the recovered DNA, homogenized, and DNA precipitated a second time as before to assess concentration and purity. A final purification step using 1.5x volumes of in-house magnetic beads GE Sera-Mag Carboxylate Modified Speed Beads) was performed according to the KAPA Pure Beads protocol (Roche, Basel, Switzerland). DNA concentration and purity were measured as described above.

### Bead Beating Optimization

Bead beating optimization was performed for three parameters: duration of bead beating, size of beads, and the bead volume. *C. neoformans* cells from overnight cultures were prepared as described above and 10^8^ cell/0.5 mL were aliquoted in threaded 2.0 mL cryotubes with 500 nm glass beads and beat for 8 cycles (1 minute on, two minutes off), with a 10 uL aliquot assessed for cell lysis following each cycle. For bead size determination, cryotubes containing *C. neoformans* cells were bead beat for 5 cycles containing 500 mg of varying bead sizes (100 nm, 500 nm, or 1000 nm), no beads, or glass shards. The glass shards were made by pulverizing 1000 nm beads into jagged, smaller pieces. For bead volume optimization, bead aliquots of 0 to 800 mg of 500 nm beads were added to *C. neoformans* lysis tubes and bead beat for 5 cycles.

Cell lysis was then inspected to quantify intact and lysed cells.

### Imaging of cell lysis

*C. neoformans* cells from overnight cultures were subjected to bead beating under a variety of conditions for optimization of cell lysis. Ten uL from each lysis sample was mixed thoroughly with 10 uL of 0.4% trypan blue solution (Thermo Fisher Scientific, Waltham, MA) and immediately visualized. Three images from each biological replicate were captured using a Leica DM 750 microscope (Leica Biosystems, Wetzlar, Germany) mounted with a Leica MCD170 HD Camera at 40x magnification. Images were processed using Leica Application Suite 4.12.0 software (Leica Biosystems, Wetzlar, Germany). Quantification of lysed cells was determined by enumeration of the cells displaying colorimetric differences between intact (clear) and lysed (blue) cells in each field of view.

### Digestion of methylated bacterial DNA

To assess digestion of 6-methyladenosine DNA, 1 ug of DNA from *E.coli*, *C.neoformans*, or an equivalent mixture of each species was digested with 1 U DpnI (NEB, Ipswitch, MA) for 3 hours at 37°C. DNA digestion was visualized on 2% TAE agarose gel.

### Magnetic bead purification assays

To assess which magnetic bead concentration optimal, three concentrations were tested: 0.7x, 0.5x, and 0.4x. A mixture of *E.coli* and *C.neoformans* digested DNA was used. Briefly, magnetic beads (GE Sera-Mag Carboxylate Modified Speed Beads) were incubated with the DNA according to the KAPA2G magnetic beads protocol and eluted in 100 uL of water.

### Mock community engineering and DNA extraction

A mock community was engineered by inoculating synthetic stool with 10 species: *C.albicans, C.neoformans, S.cerevisiae, L.major, T.brucei, C.difficile, E.coli, L.reuteri, S.typhimurium,* and *S.aureus*. For mock communities with a 1:1 ratio of bacteria to eukaryotes, still was seeded with 1 x 10^8^ cells of each species prepared as described above for *C. neoformans*. For mock communities with a 99:1 ratio of bacteria to eukaryotes, 1 x 10^10^ bacteria and 1 x 10^8^ eukaryotic cells were seeded in synthetic stool. Cell numbers were determined by hemacytometer counts for eukaryotes and OD_600_ for bacterial cells. The mock communities were then frozen at -20°C. Mock communities were extracted by adding 500 uL of inoculated synthetic stool to 500 uL of TENT buffer and 0.6 mg of 500 nm beads in lysis tubes. The samples were mechanically lysed for 5 cycles (1 minute on, 2 minutes off). DNA was purified by phenol:chloroform extraction and magnetic bead isolation as described above and resuspended in a final volume of 100 uL of water.

10 ng/uL of DNA was sheared in a Bioruptor Pico Diagenode to manufactures specifications. 20 uL of sheared DNA was then digested with 1 U of DpnI in cutsmart buffer (NEB) for 3 hours at 37°C. The reaction was stopped using 0.7x magnetic beads with a final volume of 100 uL. The effects of enrichment were measured with qPCR.

### qPCR of species abundance

All qPCR reactions were performed using primers in Table 1. Primers were designed to uniquely identify single species based on identification of hallmark single copy genes within the target organism and BLASTing the gene against the 9 other organisms in the mixed community using Ref-seq genomes to insure a lack of clear homology to other organisms. Primers were constructed using GenScript Online PCR Primers Designs Tool (https://www.genscript.com/tools/pcr-primers-designer). Primers produced a 200 base pair (bp) amplicon from these diagnostic loci. qPCR was performed from extracted DNA using an Applied Biosystems QuantStudio 5 Real-Time PCR System with PowerUp SYBR Green Master Mix (Applied Biosystems, Foster City, CA). Locus abundance was determined using a 10-fold dilution series of known DNA concentrations and generation of a standard curve equation for each organism in the mock community.

### Figures and Statistics

Each experiment was performed with at least three biological replicates and three technical replicates. GraphPad Prism 8 was used to record and analyze data with the exception of

Pearson’s correlation and linear regression models test was performed using Stata version IC.

### Ethics approval and consent to participate

Not applicable. **Consent for publication** Not applicable

### Availability of data and materials

All data generated or analysed during this study are included in this published article [and its supplementary information files].

## Authors’ contributions

M.Z.A. and A.L.C. conceived and designed experiments. A.L.C., L.M., and C.R. performed experiments and analyzed data. A.L.C. and M.Z.A. wrote the manuscript and all authors approved the final version. M.Z.A. supervised this work.

## Acknowledgements

The authors would like to thank the Anderson and Dr. Chad Rappleye labs for helpful discussion and feedback during development of this method. In particular, we would like to thank Leah Anderson for her support with reagents. We also need to thank the labs of Dr. Juan Alfonso, Dr. Vanessa Hale, and Dr. Abhay Satoskar for providing the *T. cruzi*, *C. dificile*, and *L. donovani* microbes, respectively, for the defined microbial communities used within this study.

---

Table 1. Species and species-specific primers used in this study.

Figure S1. Single copy loci used for microbial species quantification. The loci used to determine abundance of all ten species within synthetic microbial communities are depicted. The full locus is drawn and marked by the coding sequence coordinates. Gene directionality is indicated as the arrowhead marking the 3’ end of the coding sequence. Potential DpnI recognition sites (5’ GATC 3’) are indicated with black ticks. The magenta bar denotes the amplified region from qPCR.

